# Interpreting Polygenic Score Effects in Sibling Analysis

**DOI:** 10.1101/2021.07.16.452740

**Authors:** Jason Fletcher, Yuchang Wu, Tianchang Li, Qiongshi Lu

**Affiliations:** University of Wisconsin-Madison; University of Wisconsin

## Abstract

Researchers often claim that sibling analysis can be used to separate causal genetic effects from the assortment of biases that contaminate most downstream genetic studies. Indeed, typical results from sibling models show large (>50%) attenuations in the associations between polygenic scores and phenotypes compared to non-sibling models, consistent with researchers’ expectations about bias reduction. This paper explores these expectations by using family (quad) data and simulations that include indirect genetic effect processes and evaluates the ability of sibling models to uncover direct genetic effects. We find that sibling models, in general, fail to uncover direct genetic effects; indeed, these models have both upward and downward biases that are difficult to sign in typical data. When genetic nurture effects exist, sibling models create “measurement error” that attenuate associations between polygenic scores and phenotypes. As the correlation between direct and indirect effect changes, this bias can increase or decrease. Our findings suggest that interpreting results from sibling analysis aimed at uncovering direct genetic effects should be treated with caution.

## Introduction

Due to the high correlation in genetic measurement between offspring and parents, it is difficult to separate “direct” genetic effects of offspring genotype on phenotype with “indirect” genetic effects of parental genotype on offspring phenotype. This issue was demonstrated empirically by Kong et al. (2018), who showed that associations between non-transmitted parental alleles and offspring phenotype explained approximately 30% of the r-squared of the offspring polygenic scores (PGS) on offspring phenotype in the case of educational attainment.

Researchers have since made the conjecture that sibling models can solve this and other problems, since biological siblings share the same parents and therefore may share the same indirect genetic effects[1]. Indeed, many subsequent analyses have demonstrated important reductions in the estimated associations between PGS and phenotypes after controlling for sibling fixed effects, often on the order of 50% (Trejo and Domingue 2018, Selzam et al. 2019). This theory and evidence have increased researchers’ confidence in interpreting sibling models as producing causal (direct) genetic effects—both in upstream GWAS analysis (Howe et al. 2021) and in downstream PGS analysis (Belsky et al. 2018).

While intuitive, these interpretations have not been subject to theoretical and empirical scrutiny. We use family data (quads) combined with simulation evidence to show that this intuition relies on very simple models of genotype-to-phenotype associations, where direct and indirect genetic effects are separable (and thus differenced out in sibling models). While researchers have suggested that using a sibling model would purge the indirect effects, we show that, even in our simplified cases, researchers have failed to recognize the dual role of indirect genetic effects as a confounder that produces both positive and negative bias on the estimated effects of offspring PGS. We build the intuition for the negative bias by considering a scenario where all SNP effects are causal and there are no indirect genetic effects and show that sibling analysis leads to attenuated estimates of PGS effects. Essentially, the attenuation stems from sibling analysis’ elimination of both confounding and true genetic effects. We then extend this scenario to allow correlation between indirect and direct genetic effects to show that a combination of biases in general can be negative or positive and depend on the strength of indirect effects, the correlation between indirect and direct genetic effects, and also the extent to which sibling’s environments are correlated. In summary, even our simple data generating process shows that sibling analysis is unlikely to produce accurate estimates of direct effects and, worse, does not suggest clear correctives or bounding exercises that are effective.

## Result

The detailed simulation procedures are described in the Materials and methods section. Briefly, we draw the direct and indirect effect sizes for each SNP as well as the environmental effect for each individual following a specific set of parameters in each scenario. Then, we regressed the phenotype constructed by adding up the components above on the theoretical PGS estimation that can be obtained from outcome of genome-wide association studies (GWAS). We compared either the regression coefficient or R^2^ of both between- and within-family regression to imitate real PGS analysis and investigate the impact of changes of parameters in the data generating process on the performance of sibling PGS analysis.

### The influence of the correlation of environmental effects between siblings

We performed simulations under the simple scenario where the phenotype has no indirect genetic effect contribution (*σ*_*i*_ = 0). In this case, from formula (10, 11) in the method section, we can see that sibling PGS models are not guaranteed to estimate the variance component of direct effects even when genetic nurture is absent. This is because the elimination of family effects that occur in sibling models also eliminate true direct genetic effects. Our formulas show that the elimination of true genetic effects can be offset in the (unlikely) case where siblings have a correlation of 0.5 in their environmental effects (*ρ*_*e*_= 0.5); in this case, the correlation in both environmental and genetic effects are the same. In general, when siblings have a correlation in their environmental effects higher than 0.5, the R^2^ of sibling PGS model is overestimated; when this correlation drops below 0.5, the R^2^ of sibling PGS model underestimates (Fig 1). Since the environmental effect is assumed to be independent of genetic components in this study, this factor does not interact with other potential factors. Therefore, to focus on the influence of other parameters (and not *ρ*_*e*_), we will assume that siblings have a correlation of 0.5 in their environmental effects throughout the rest of the simulations. Results under correlations of 0 and 1 are included in the supplementary table.

**Fig 1.**
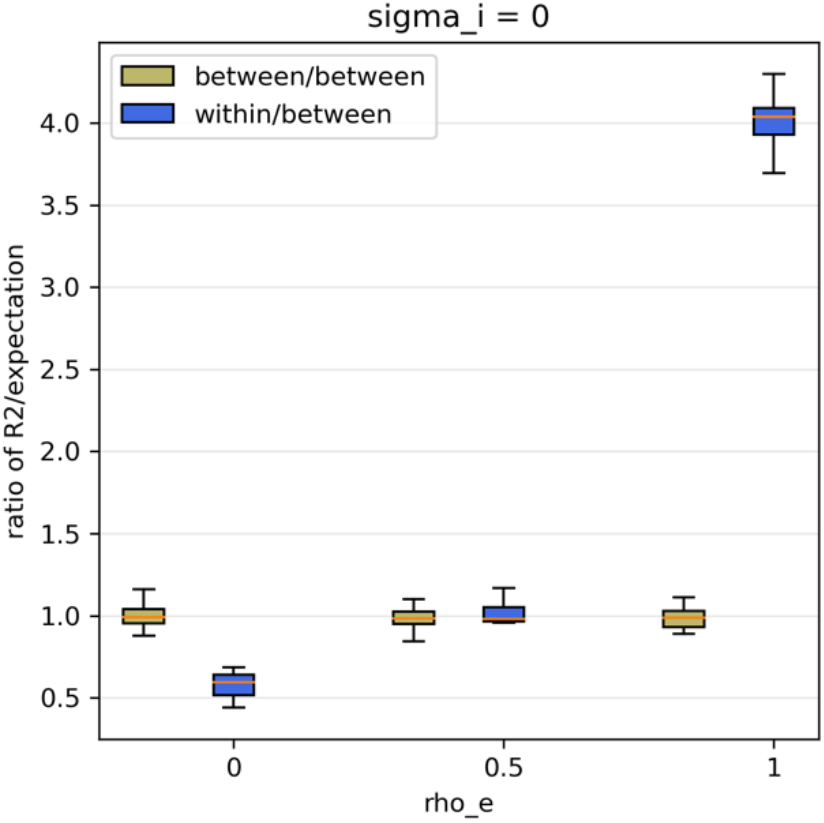
Comparison of the ratio of PGS regression R^2^ and true direct effect variance component under different values of sibling environmental correlation. Two estimates of interest included are the R^2^ obtained from between-family (yellow) and within-family regressions (blue) when indirect effect is absent. Boxplots display the ratio of the R^2^ estimated on 15 simulation repetitions over the designed population-level direct effect variance component on y-axis versus various values of sibling environmental correlation from 0 to 1 on x-axis. The ratio for between-family R^2^ remain at 1 while the ratio for within-family R^2^ fluctuate except when the sibling environmental correlation equals 0.5 under our assumptions. This suggests that sibling environmental correlation is an additional factor that impacts the performance of sibling PGS analysis even with no indirect genetic effects.

### The influence of indirect genetic effects

We performed simulations with the variance of direct genetic effects normalized to 1, the variance of the environmental effect is assumed to be three times the variance of the direct genetic effect, and we assume no correlation between direct and indirect genetic effects to concentrate on the influence of indirect genetic effects. With we increase the contribution of indirect genetic effects, we found that sibling PGS models produce estimates of the direct effect variance component that is attenuated compared to the estimates for the population (i.e. non-sibling) PGS models (Fig 2). When indirect genetic effects are absent, the sibling PGS model estimates accurately (see formula (10, 11)) (recall, we assume *ρ*_*e*_ = 0.5 so that we can focus only on other biases from indirect genetic effects). Then, as the indirect genetic effect is introduced, the downward bias of the sibling model becomes larger. This is consistent with empirical results from other work cited above. However, since the ratio of these estimated R^2^ values compared with the expected direct effect component variance is smaller than 1, our results demonstrate that the sibling PGS model does not accurately recover the direct effect heritability. The figure also shows that the population model cannot recover direct effects, and that the estimated effects are biased upward (as is commonly understood).

**Fig 2.**
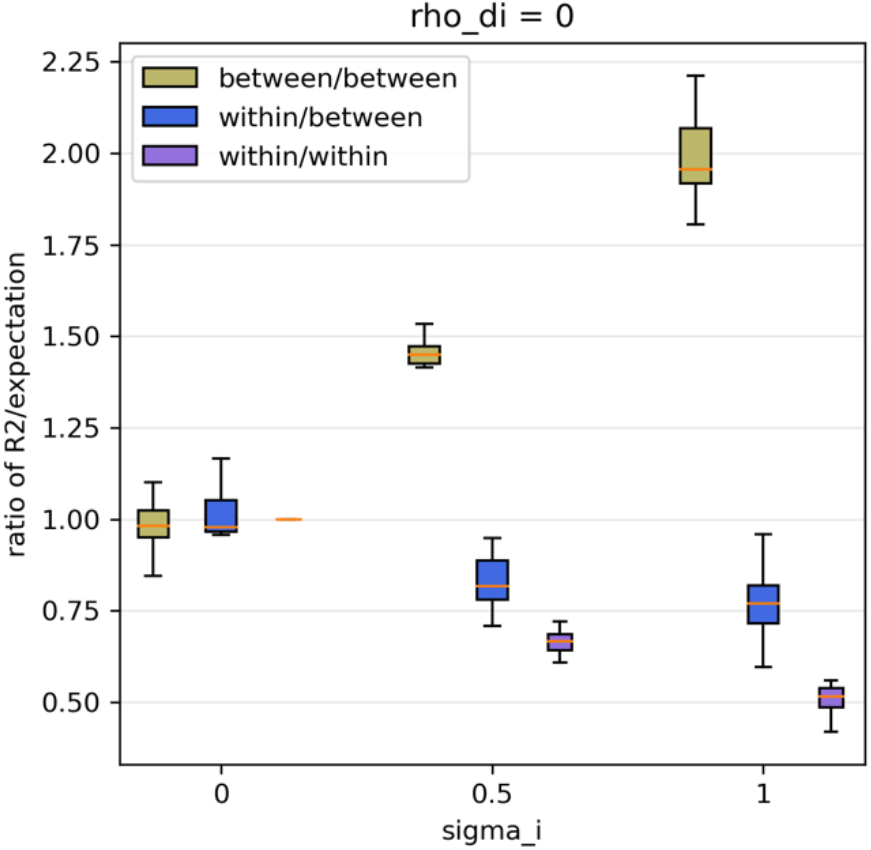
Comparison of the ratio of PGS regression R^2^ compared with the true direct effect component variance under different values of indirect genetic effect variance component. Two estimates of interest included are the R^2^ obtained from between-family and within-family regression when direct and indirect effects are independent. Boxplots display the ratio of these two R^2^ estimated on 15 simulation repetitions over the designed population-level direct effect component variance (yellow and blue) and the ratio of within-family R^2^ over expected within-family R^2^ estimated with only direct effect (purple) on y-axis versus various values of indirect effect component variance from 0 to 1 on x-axis. The downward trend from 1 of blue and purple boxes as indirect effect component variance increases suggests that indirect effect only biases the sibling R^2^ downward as opposite to biasing the between-family R^2^ upwards as indicated by yellow boxes.

We now consider the regression that sibling PGS model (i.e. sibling difference model) performs, 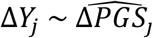. When taking the difference in sibling phenotypes, the genetic nurture effect cancels out since full siblings share the same parents (formula (6)). Whereas in estimated PGS, the transmitted genetic nurture remains different between siblings (formula (7)). To better understand the impact of indirect genetic effects on a sibling PGS analysis, we plot the ratio of sibling model R^2^ estimated with 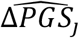 compared with the R^2^ estimated with the sibling difference in direct effect component (formula (12)). This compares the regular sibling model estimated R^2^ with the R^2^ of true direct effects. Note in the third box in each group, we see results below 1, we can see that the indirect effect reduces the regression R^2^ more as its contribution to the phenotype increases.

### The influence of correlation between direct and indirect effects

As above, we constructed phenotypes with the variance of direct effect normalized to 1, the variance of indirect genetic effect as either a half or equal to the variance of the direct genetic effect, and the variance of environmental effect as three times the variance of the direct genetic effect. We now relax the assumption from above that the correlation between indirect and direct genetic effects is zero. Now, we varied the correlation between direct and indirect effects between −1 and 1, and we found that sibling PGS models, in general, produce estimates of the variance component of direct effect that are smaller than estimates of population PGS models (Fig 3). We note that previous literature viewed reductions in estimated direct effect contribution using sibling models as evidence that these models were eliminating confounds (such as indirect genetic effects), whereas our results show that sibling models actually underestimate direct genetic effects in many scenarios.

**Fig 3.**
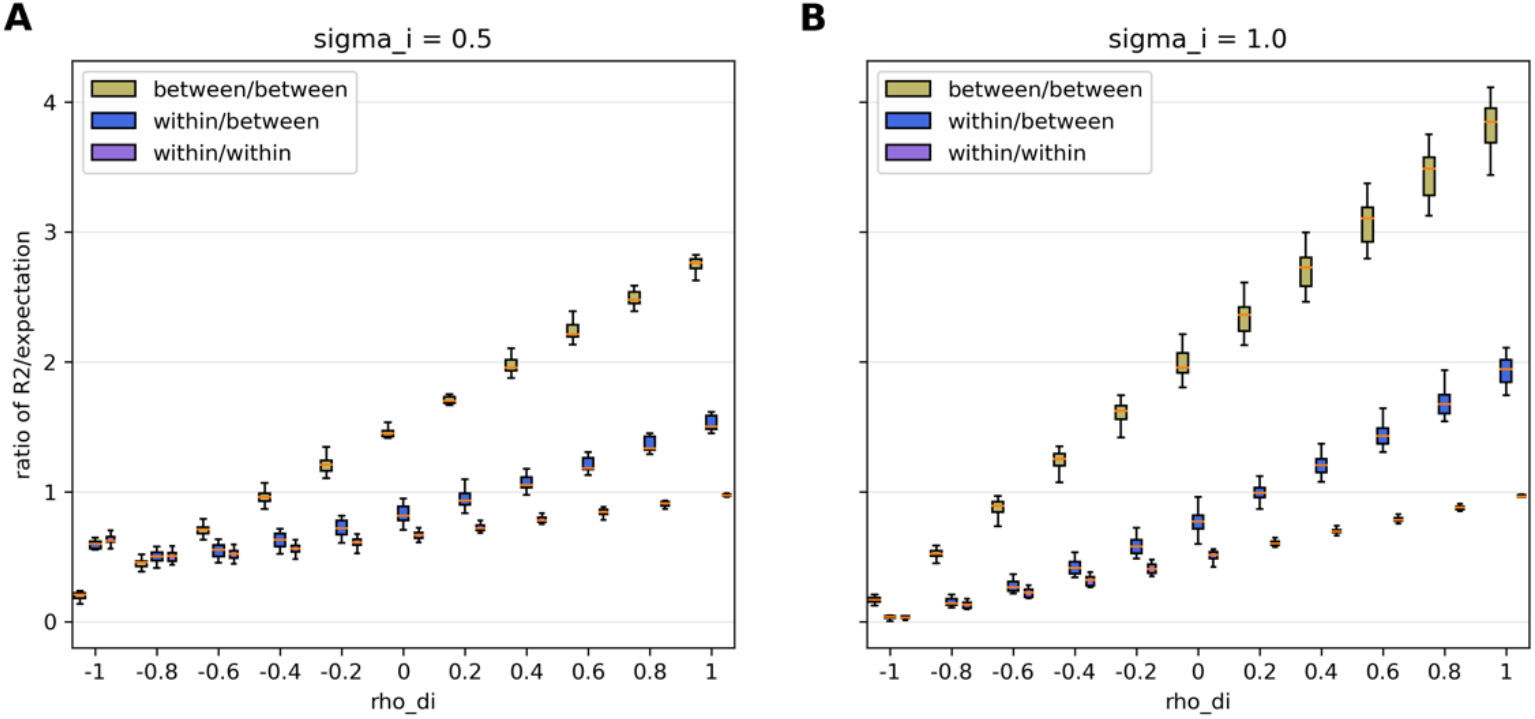
Comparison of the ratio of PGS regression R^2^ over true direct effect component variance under different values of direct and indirect effect correlation. Two estimates of interest included are the R^2^ obtained from between-family and within-family regression when indirect effect component variance equals to 0.5 (A) and 1 (B) respectively. Boxplots display the ratio of these two R^2^ estimated on 15 simulation repetitions over the designed population-level direct effect component variance (yellow and blue) and the ratio of within-family R^2^ over expected within-family R^2^ estimated with only direct effect (purple) on y-axis versus various values of direct and indirect effect correlation from −1 to 1 on x-axis. As the correlation deviates from 0, the PGS regression R^2^ is either inflated or deflated.

For each of the two settings that fix the variances of direct and indirect genetic effects, we show that increasing the correlation between direct and indirect effects increases the ratio of sibling PGS estimates and the direct effect component variance from below 1 (underestimate) to above 1 (overestimate). R^2^ can be viewed as a function of the correlation between direct and indirect effects given specific values of other parameters, such as the variance of indirect effect. So, the ratio of sibling PGS estimate and the direct effect variance component defined on population does not change linearly (formula (10), Appendix 3). Whether we allow indirect genetic effects to be modest (0.5) or large (1) and also allow correlations between the direct and indirect genetic effects, we find that it is rare that sibling analysis will accurately estimate direct genetic effects (i.e the horizontal line at 1, where the estimated R^2^ is equal to the expected R^2^). The results also suggest that the estimated R^2^ can be up to twice the size of the true R^2^ or less than half the size of the true R^2^, depending on the data generating process. We also note again that we have assumed in these analyses that the environmental effects are correlated at 0.5 between siblings. Otherwise, these results would “shift down”, as we show in Fig 1.

### The performance of PGS regression coefficients

As another important measurement of PGS models’ performance, regression coefficients have also drawn attention in the past. One study showed that between-family PGS regression would yield coefficients that are biased upwards and within-family PGS regression would yield coefficients that bias downwards (Trejo & Domingue, 2018). To verify this in our framework, we estimated the regression coefficients under our setup on a wider value range of rho. We also noted a divergence between our framework and the previous one in terms of the standardization of PGS. Our framework does not standardize the PGS variable in the estimation, so that the between-family regression will estimate a coefficient of 1, which equals the theoretical value when direct effect variance component is accurately recovered (Fig 4). For the sibling PGS model, with unstandardized PGS estimates, regression coefficients could be over- or under-estimated depending on the ratio of indirect and direct effect variance and the value of their correlation. Overall, the larger the ratio is, the more the sibling regression coefficients converge to a downwardly biased value. The larger the correlation is, the more the sibling regression coefficients are biased downwards (Fig 4). We also confirmed these with our derivation and rewrite Trejo and Domingue’s derivation without standardization of PGS (Appendix 5).

**Fig 4.**
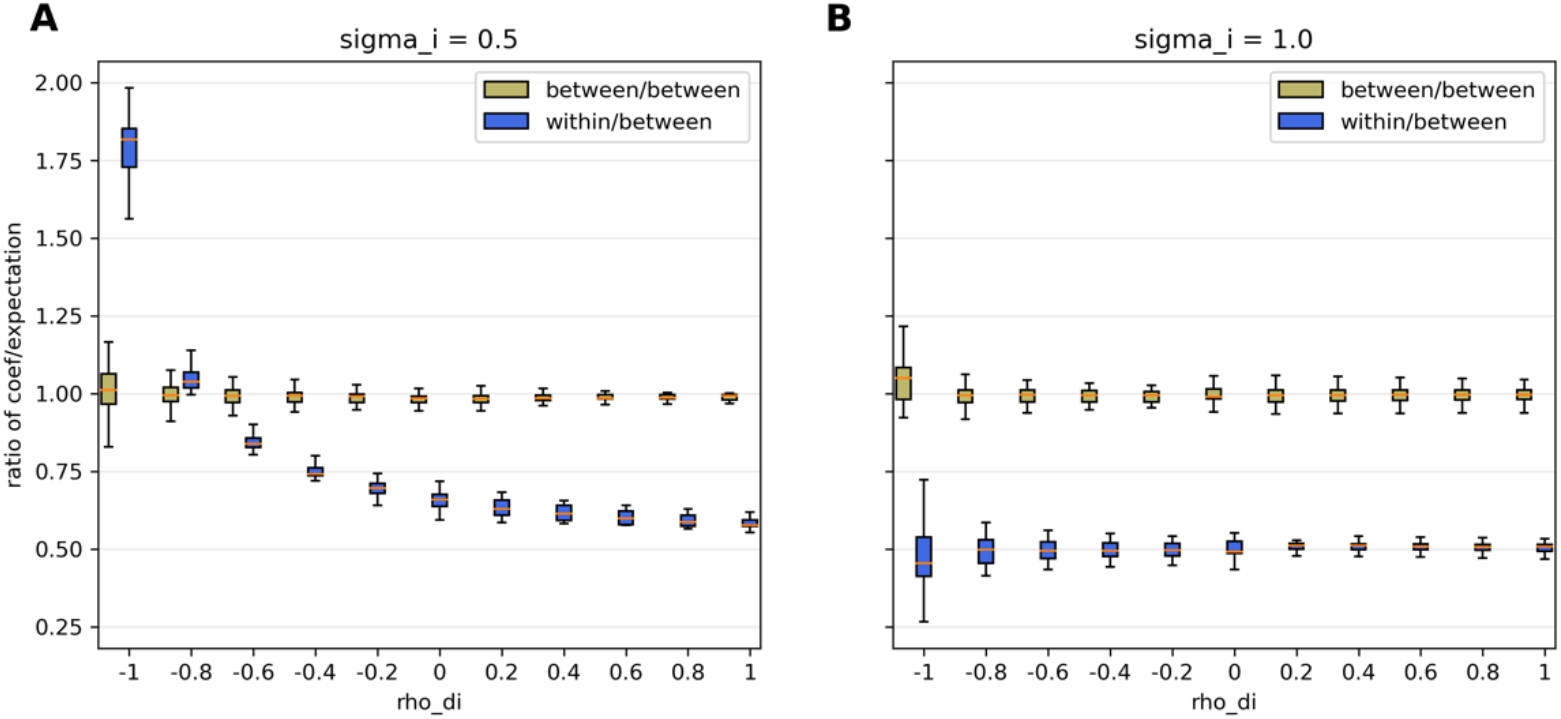
Comparison of the ratio of PGS regression coefficients over true direct effect component variance under different values of direct and indirect effect correlation. Two estimates of interest included are the regression coefficients obtained from between-family (yellow) and within-family (blue) regression when indirect effect variance component equals to 0.5 (A) and 1 (B) respectively. Boxplots display the ratio of these two coefficients estimated on 15 simulation repetitions over the expected coefficient for direct effect component, 1, on y-axis versus various values of direct and indirect effect correlation from −1 to 1 on x-axis. They suggest that between-family regression coefficient is always estimated to be 1 regardless of the direct and indirect effect correlation whereas within-family regression coefficients are mostly biased downwards but also depend on the indirect effect variance component.

## Discussion

To examine the performance of the estimated R^2^ of sibling PGS analysis in recovering the direct genetic effect variance component from data generating process that includes both direct and indirect genetic effects, we performed our analysis on simulated phenotypes based on genotypic data of quads in the SPARK cohort. Our analytical results demonstrated that sibling PGS analysis generally does not yield R^2^ that accurately reflects the direct effect variance component.

In the simplified scenario where a phenotype is not impacted by indirect genetic effects, sibling PGS analysis could yield R^2^ biases either upward or downward depending on the environmental correlation between siblings. In this study, we assumed a genotypic correlation of 0.5 between siblings which set a scale for the variance of the difference between their estimated PGS. More importantly, when taking the difference in sibling phenotypes, the indirect effect that is shared between siblings is eliminated, leaving only the difference in direct effect PGS and difference in environmental effects. However, PGS constructed from GWAS estimates will, in most cases, contain both direct and indirect effects. When taking a difference between siblings, direct genetic effect and transmitted genetic nurture effects remain in the beta weights that are used to construct the PGS in downstream analysis. Given these issues, three aspects of the sibling PGS model are found to generate biases.

Firstly, based on the composition of sibling phenotype difference, sibling PGS regression only retains the proportion explained in direct genetic effect and environmental effect differences. Essentially, when the phenotype is also affected by indirect genetic effects, the total variance of the phenotype is reduced when using sibling analysis compared to the variance defined in population-based PGS regression. Even with accurate direct effect PGS, sibling PGS would still fail to fully recover the direct effect variance component for the population. Then, in order to examine the impact of other factors, we turned to comparing the sibling R^2^ estimates with the theoretical sibling R^2^ when regressing phenotype difference on true direct PGS difference.

We found that, as the contribution of indirect genetic effects increases from 0, the ratio of estimated sibling R^2^ and the theoretical sibling R^2^ of direct effect PGS continues to decline from its target value of 1. This means the indirect effect component is similar to “measurement error” in this case, attenuating the direct effect estimates. Additionally, when the contribution of direct, indirect, and environmental effects is held fixed, changes in the correlation between direct and indirect effect will also lead to bias in sibling R^2^. As the correlation reduces from 1 to −1, the estimated R^2^ is increasingly biased downwards. We label this as an “LD-like” relationship between direct and indirect genetic components, which survives sibling differencing or sibling fixed effects analysis. When compared with the direct effect variance component defined at the population level, this could lead to either downward or upward bias. Similarly, we make slight adjustments on the previous results from Trejo and Domingue on the bias of regression coefficients obtained from sibling PGS analysis. Our conclusion shows that sibling analysis would still be biased upwards or downwards depending on the combinations of variances of direct genetic effects, indirect genetic effects, and their correlation.

It is important to note that our results are from a “simple” data generating processes, where we assumed no assortative mating, no gene-environment interaction or correlation, and sibling genetic correlations of exactly 0.5. Adding these other elements to the framework will further complicate evaluating the performance of the sibling analysis, but, we suspect, will lead to additional biases rather than fewer. Thus, our view of the results from a relatively simple framework is that sibling analysis, coupled with conventional PGS, can rarely uncover a key target—direct genetic effects. Solving this issue will rely on dissecting each individual variant’s direct and indirect effects and calculating respective PGS for direct and indirect genetic components, possibly through sibling GWAS and multi-generational analysis (Howe et al., 2021; Wu et al., 2021).

## Materials and methods

### Data

We leverage family-based genetic data from quads (2 parents, 2 children) in the SPARK (Simons Foundation Powering Autism Research for Knowledge) study (Feliciano et al 2018) in order to seed the model with realistic genetic information. Specifically, we obtained 7,026,791 SNPs from 1813 families with two parents and two full siblings. Following previous work (Huang et al. 2021), we filtered out single nucleotide polymorphisms (SNPs) with a minor allele frequency less than 1%, with an imputation quality score less than 0.8, that are duplicated, or strand-ambiguous. Then we pruned the SNPs with linkage disequilibrium (LD) with a pairwise r^2^ higher 0.1. From the remaining 127,310 SNPs, we randomly picked 10,000 SNPs as causal variants and simulated phenotypes based on them.

### Model specifications

We assume a data generating process for the phenotype Yij that includes both direct and indirect genetic effects. Our model follows Kong et al. (2018) by assuming additive separable direct (*β*_*dir,k*_) and indirect (*β*_*ind,k*_) genetic effects that are drawn from a bivariate normal distribution with a correlation parameter. The indirect genetic effect behaves as a family fixed effect that is shared between siblings. We denote the genotype of the i^th^ child/sibling in the j^th^ family as Gij; the genotype of mother and father in the j^th^ family as G_m,j_ and G_p,j_ respectively. We can write the model as

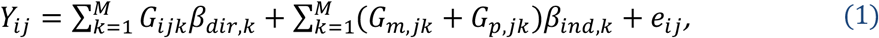

where e_ij_ is the environmental residual. We assume equal indirect paternal and indirect maternal effect, both being *β*_*ind,k*_. We note that this parameter can also be viewed as the average indirect parental effect if maternal and paternal effects are in fact unequal (Wu et al. 2021). We also assume that the direct and indirect effect sizes of the k^th^ SNP on the phenotypes follow

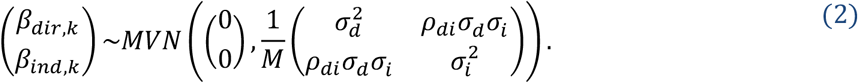

where 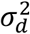 represents the variance of the direct effect component of the phenotype; 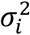 represents the variance of the indirect effect component of the phenotype; and *ρ*_*di*_ represents the correlation between direct and indirect genetic effect sizes; M denotes the number of causal SNPs with direct effect size *β*_*dir,k*_ and indirect effect size *β*_*ind,k*_. For families whose genotypes are used in the simulation, we assume that the environmental effects for the 2 children in the j^th^ family also follow a bivariate normal distribution

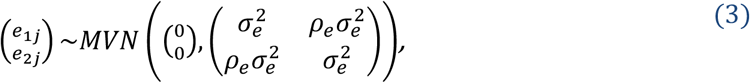

where 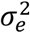 represents the variance of environmental effects shared between 2 siblings; and *ρ*_*e*_ represents the environmental correlation between 2 siblings. We further assume all genotypes involved are standardized.

Formula (1) can be rearranged to separate the effects of transmitted (G_ij_) and non-transmitted (N_ij_) alleles as

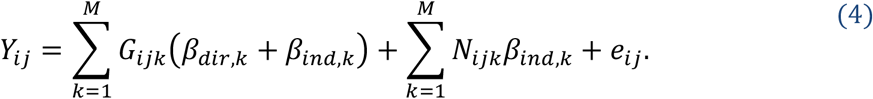

We assumed transmitted and non-transmitted alleles to be independent (supported by genotypic data, Appendix 1). It is clear to see from the rearrangement that a GWAS on phenotype will capture both the true direct and indirect genetic effects. Following the conventions in the literature, we constructed the downstream PGS estimation with the theoretical GWAS estimated allelic weights 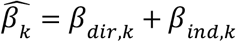, assuming all causal SNPs are accurately estimated, which we denote as 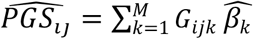 (Lee et al. 2018). We obtained between-family PGS regression coefficients *γ* _*OLS*_ and r-squared *R*^2^ _*OLS*_ by regressing the phenotype of one sibling from each family on their estimated PGS as

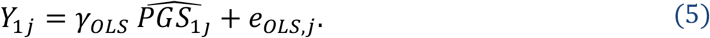

Derivations on the theoretical regression coefficient and r-squared for between-family analysis are included in the Appendix 2. For sibling analysis, we took the difference in the phenotype between two siblings in a family as the within-family outcome

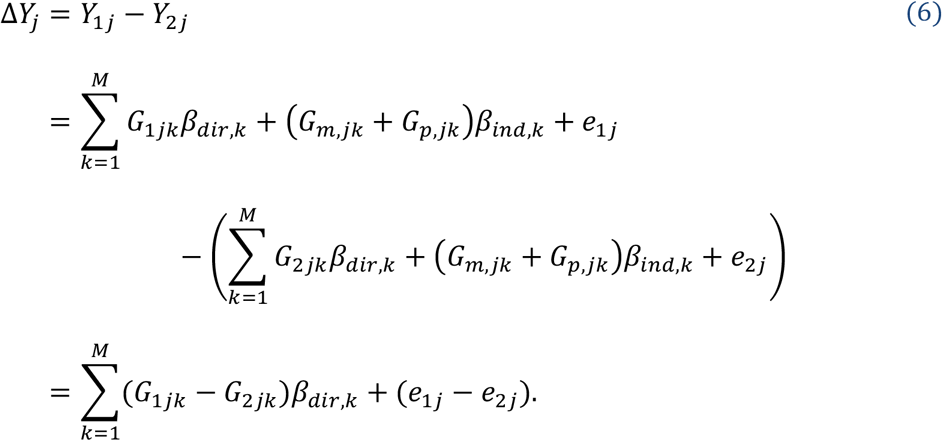

The shared indirect effect is eliminated between siblings. We took the difference in the estimated PGS between two siblings in a family as the within-family predictor

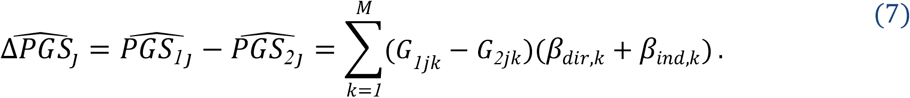

Then we obtained within-family PGS regression coefficients *γ* _Δ_ and r-squared *R*^2^ _Δ_ by regressing the difference in the phenotype on the difference in the estimated PGS as

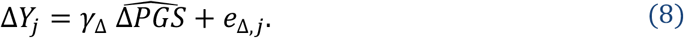

When we assume siblings from the same families have a correlation of 0.5 in their genotypes, *G*__1jk__ and *G*__2jk__ (also supported by the genotypic data we use for simulation, details in Appendix 1), the within-family regression coefficients and r-squared can be derived as

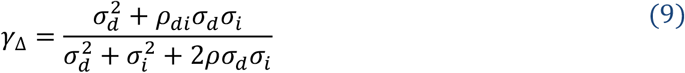

and

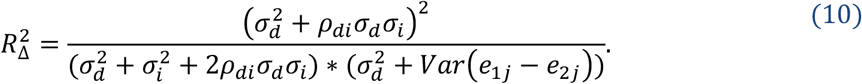

To quantify the performance of both analyses on recovering the direct effect component, we compared the outcome regression coefficient with 1 (as direct effect component in model (2) takes a coefficient of 1) and r-squared with the proportion of direct effect variance component defined on population base

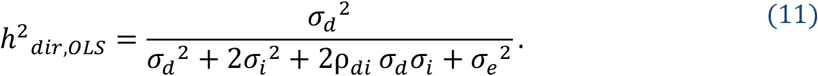

We also compared the outcome r-squared of sibling analysis with the proportion of direct effect variance component defined on sibling differences which allowed us to better understand the impact of the change of parameters on sibling analysis alone. That is, we performed regression

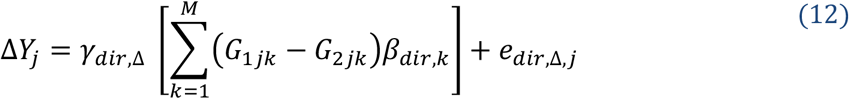

and obtained

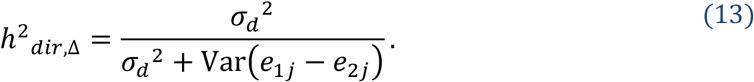

(Appendix 4)

### Simulation

We generated direct effect and indirect effect allelic weights for each offspring from a normal distribution from each combination of parameters and apply them to offspring’s standardized genotypes. We also generated environmental effects for each offspring from a normal distribution following the parameters in each setting. By adding these components up for each offspring, we obtain their phenotypes.

#### Setting 1

From the derivations of our estimates above, we found that even in the simplest scenarios with unbiased GWAS effect sizes and genetic nurture absent, sibling analysis does not accurately estimate the variance component of direct effect. As a special case of formula (11), here we have the population-based direct effect variance component defined as

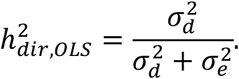

However, R^2^ from sibling analysis is expected to be

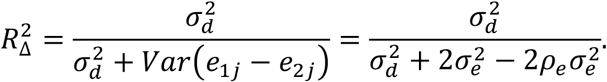

Comparing these two formulas, one can see that the difference between them depends on the correlation between the environmental effect of siblings, *ρ*_*e*_. Only when *ρ*_*e*_ = 0.5, these two quantities equal. Therefore, we designed a setting where we kept the variance of direct genetic effect constant and set indirect genetic effect to be 0. Then, the correlation between direct and indirect effect is also 0. We also set the variance of the environmental residual and set the correlation between siblings’ environmental residual, *ρ*_*e*_, to 0, 0.5, or 1. Thus, a total of 3 scenarios were examined in setting 1.

#### Setting 2

We kept the variance of direct genetic effect constant (normalized to 1) and varied the indirect genetic effect and the correlation between direct and indirect genetic effect to evaluate the influence of each factor on the sibling analysis R^2^ when the other was held constant. Specifically, we set the variance of indirect effect to be either 0, 0.5, or 1. For each variance of indirect effect, we varied the correlation between direct and indirect effect from −1 to 1 by a step of 0.2. We also set the variance of environmental residual to be 3. A total of 33 scenarios (3 variance of indirect effect x 11 correlation between direct and indirect effect) were examined in setting 2.

## Acknowledgements

The authors gratefully acknowledge use of the facilities of the Center for Demography of Health and Aging at the University of Wisconsin-Madison, funded by NIA Center Grant P30 AG017266. We thank members of the Social Genomics Working Group at University of Wisconsin for helpful comments. We are grateful to all the families participating in the Simons Foundation Powering Autism Research for Knowledge (SPARK) study.

## Footnotes

[1] The more general claim of the usefulness of siblings and the associated “genetic lottery” proceeds the results of indirect genetic effects (see Fletcher and Lehrer 2009, 2011)

