## Supplemental Appendix for "Interpreting Polygenic Score Effects in Sibling Analysis"

### 1. Genotype standardization and assumptions

We assume all genotypes are standardized. That is,

$$E(G_{1jk}) = E(G_{2jk}) = E(G_{m,jk}) = E(G_{p,jk}) = 0;$$
$$Var(G_{1jk}) = Var(G_{2jk}) = Var(G_{m,jk}) = Var(G_{p,jk}) = 1.$$

The transmitted and non-transmitted alleles of each child in the SPARK dataset have an average correlation of -0.012 ($1^{st}$ quantile = -0.029, $3^{rd}$ quantile = 0.006). The PGS created from transmitted and non-transmitted alleles of each child has a correlation of =-0.056. Based on these empirical results, $G_{ij}$ and $N_{ij}$ can be reasonably assumed to be independent. Then, we have

$$E(N_{1jk}) = E(N_{2jk}) = E(G_{m,jk} + G_{p,jk} - G_{1jk}) = E(G_{m,jk} + G_{p,jk} - G_{2jk}) = 0;$$
$$Var(N_{1jk}) = Var(N_{2jk}) = Var(G_{m,jk} + G_{p,jk} - G_{1jk}) = Var(G_{m,jk} + G_{p,jk} - G_{2jk}) = 1$$

The genotypes of two siblings in each family have an average correlation of 0.4732 ($1^{st}$ quantile = 0.4679, $3^{rd}$ quantile = 0.5154), which supports the assumption that the correlation of $G_{1jk}$ and $G_{2jk}$ is 0.5. Then, we have

$$Var(G_{1jk} - G_{2jk}) = 1 + 1 - 2 * 0.5 = 1$$

### 2. Between-family PGS regression estimates

Regressing the phenotype of one sibling from each family (i.e., independent samples) on their PGS, i.e.,

$$Y_{1j} = \gamma_{OLS}\widehat{PGS_{1j}} + e_{OLS,j},$$

we expect to obtain regression coefficient and $R^2$ as follows

$$\begin{aligned}\gamma_{OLS} &= \frac{Cov(\widehat{PGS_{1j}}, Y_{1j})}{Var(\widehat{PGS_{1j}})} \\ &= \frac{Cov\left(\sum_{k=1}^{M} G_{1jk}(\beta_{dir,k} + \beta_{ind,k}), \sum_{k=1}^{M} G_{1jk}(\beta_{dir,k} + \beta_{ind,k}) + \sum_{k=1}^{M} N_{1jk}\beta_{ind,k} + e_{1j}\right)}{Var\left(\sum_{k=1}^{M} G_{1jk}(\beta_{dir,k} + \beta_{ind,k})\right)} \\ &= \frac{Cov\left(\sum_{k=1}^{M} G_{1jk}(\beta_{dir,k} + \beta_{ind,k}), \sum_{k=1}^{M} G_{1jk}(\beta_{dir,k} + \beta_{ind,k})\right)}{Var\left(\sum_{k=1}^{M} G_{1jk}(\beta_{dir,k} + \beta_{ind,k})\right)} \\ &= \frac{Var\left(\sum_{k=1}^{M} G_{1jk}(\beta_{dir,k} + \beta_{ind,k})\right)}{Var\left(\sum_{k=1}^{M} G_{1jk}(\beta_{dir,k} + \beta_{ind,k})\right)} \\ &= 1;\end{aligned}$$

$$
\begin{aligned}
R^2_{OLS} &= \gamma^2_{OLS} * \frac{Var\left(\widehat{PGS_{1j}}\right)}{Var\left(Y_{1j}\right)} \\
&= \frac{Var\left(\sum_{k=1}^{M} G_{1jk}\left(\beta_{dir,k} + \beta_{ind,k}\right)\right)}{Var\left(\sum_{k=1}^{M} G_{1jk}\left(\beta_{dir,k} + \beta_{ind,k}\right) + \sum_{k=1}^{M} N_{1jk}\beta_{ind,k} + e_{1j}\right)} \\
&= \frac{Var\left(\sum_{k=1}^{M} G_{1jk}\left(\beta_{dir,k} + \beta_{ind,k}\right)\right)}{Var\left(\sum_{k=1}^{M} G_{1jk}\left(\beta_{dir,k} + \beta_{ind,k}\right)\right) + Var\left(\sum_{k=1}^{M} N_{1jk}\beta_{ind,k}\right) + Var\left(e_{1j}\right)} \\
&= \frac{M * \left(\frac{\sigma_d^2}{M} + \frac{\sigma_i^2}{M} + \frac{2\rho_{di}\sigma_d\sigma_i}{M}\right)}{M * \left(\frac{\sigma_d^2}{M} + \frac{\sigma_i^2}{M} + \frac{2\rho_{di}\sigma_d\sigma_i}{M}\right) + M * \frac{\sigma_i^2}{M} + 2\sigma_e^2 - 2\rho_e\sigma_e^2} \\
&= \frac{\sigma_d^2 + \sigma_i^2 + 2\rho_{di}\sigma_d\sigma_i}{\sigma_d^2 + 2\sigma_i^2 + 2\rho_{di}\sigma_d\sigma_i + 2\sigma_e^2 - 2\rho_e\sigma_e^2}.
\end{aligned}
$$

When the correlation of environmental effect between siblings (i.e., $\rho_e$) is 0.5, this can be simplified as

$$
R^2_{OLS} = \frac{\sigma_d^2 + \sigma_i^2 + 2\rho_{di}\sigma_d\sigma_i}{\sigma_d^2 + 2\sigma_i^2 + 2\rho_{di}\sigma_d\sigma_i + \sigma_e^2}.
$$

The ratio of it over population-level direct effect variance component $h^2{}_{dir,OLS}$ can be derived as

$$
\frac{R^2_{OLS}}{h^2{}_{dir,OLS}} = \frac{\sigma_d^2 + \sigma_i^2 + 2\rho_{di}\sigma_d\sigma_i}{\sigma_d^2 + 2\sigma_i^2 + 2\rho_{di}\sigma_d\sigma_i + \sigma_e^2} * \frac{{\sigma_d}^2 + 2{\sigma_i}^2 + 2\rho_{di}\,\sigma_d\sigma_i + {\sigma_e}^2}{{\sigma_d}^2} = \frac{\sigma_d^2 + \sigma_i^2 + 2\rho_{di}\sigma_d\sigma_i}{{\sigma_d}^2}.
$$

When the direct and indirect effects have a correlation of 0 (i.e., $\rho_{di} = 0$), this ratio equals

$$
\frac{R^2_{OLS}}{h^2{}_{dir,OLS}} = \frac{\sigma_d^2 + \sigma_i^2}{{\sigma_d}^2};
$$

and when the indirect effect is set to zero (i.e., $\sigma_i = 0$), we have

$$
\frac{R^2_{OLS}}{h^2{}_{dir,OLS}} = \frac{\sigma_d^2}{{\sigma_d}^2} = 1.
$$

### 3. Within-family PGS regression estimates

Regressing differenced sibling phenotypes on differenced sibling PGS, i.e.,

$$
Y_{1j} - Y_{2j} = \gamma_\Delta\left(\widehat{PGS_{1j}} - \widehat{PGS_{2j}}\right) + e_{\Delta,j}
$$

we expect to obtain regression coefficient

$$\gamma_\Delta = \frac{Cov(\widehat{PGS_{1J}} - \widehat{PGS_{2J}}, Y_{1j} - Y_{2j})}{Var(\widehat{PGS_{1J}} - \widehat{PGS_{2J}})}$$
$$= \frac{Cov(\sum_{k=1}^{M}(G_{1jk} - G_{2jk})(\beta_{dir,k} + \beta_{ind,k}), \sum_{k=1}^{M}(G_{1jk} - G_{2jk})\beta_{dir,k} + (e_{1j} - e_{2j}))}{Var(\sum_{k=1}^{M}(G_{1jk} - G_{2jk})(\beta_{dir,k} + \beta_{ind,k}))}$$
$$= \frac{\sum_{k=1}^{M}[Var((G_{1jk} - G_{2jk})\beta_{dir,k}) + Cov\left((G_{1jk} - G_{2jk})\beta_{ind,k}, (G_{1jk} - G_{2jk})\beta_{dir,k}\right)]}{\sum_{k=1}^{M} Var((G_{1jk} - G_{2jk})(\beta_{dir,k} + \beta_{ind,k}))}$$
$$= \frac{\sum_{k=1}^{M} 2Var(\beta_{dir,k}) - 2Cov(G_{1jk}\beta_{dir,k}, G_{2jk}\beta_{dir,k}) + \sum_{K=1}^{M} Var(G_{1jk} - G_{2jk}) * Cov(\beta_{ind,k}, \beta_{dir,k})}{\sum_{k=1}^{M} 2Var(\beta_{dir,k} + \beta_{ind,k}) - 2Cov\left(G_{1jk}(\beta_{dir,k} + \beta_{ind,k}), G_{2jk}(\beta_{dir,k} + \beta_{ind,k})\right)}$$
$$= \frac{M * \left(\frac{2\sigma_d^2}{M} - \frac{2\sigma_d^2}{M} * Cov(G_{1jk}, G_{2jk})\right) + M * \frac{\rho_{di}\sigma_d\sigma_i}{M} * \left(2 - 2Cov(G_{1jk}, G_{2jk})\right)}{M * 2\left(\frac{\sigma_d^2}{M} + \frac{\sigma_i^2}{M} + \frac{2\rho_{di}\sigma_d\sigma_i}{M} - Cov(G_{1jk}, G_{2jk}) * \left(\frac{\sigma_d^2}{M} + \frac{\sigma_i^2}{M} + \frac{2\rho_{di}\sigma_d\sigma_i}{M}\right)\right)}$$
$$= \frac{2\sigma_d^2\left(1 - Cov(G_{1jk}, G_{2jk})\right) + 2\rho_{di}\sigma_d\sigma_i * \left(1 - Cov(G_{1jk}, G_{2jk})\right)}{2(\sigma_d^2 + \sigma_i^2 + 2\rho_{di}\sigma_d\sigma_i)(1 - Cov(G_{1jk}, G_{2jk}))}.$$

and regression $R^2$

$$R_\Delta^2 = \gamma_\Delta^2 * \frac{Var(\widehat{PGS_{1J}} - \widehat{PGS_{2J}})}{Var(Y_{1j} - Y_{2j})}$$
$$= \gamma_\Delta^2 * \frac{Var(\sum_{k=1}^{M}(G_{1jk} - G_{2jk})(\beta_{dir,k} + \beta_{ind,k}))}{Var\left(\sum_{k=1}^{M}(G_{1jk} - G_{2jk})\beta_{dir,k} + (e_{1j} - e_{2j})\right)}$$
$$= \gamma_\Delta^2 \frac{\sum_{k=1}^{M} Var(G_{1jk} - G_{2jk}) * Var(\beta_{dir,k} + \beta_{ind,k})}{\sum_{k=1}^{M} 2Var(\beta_{dir,k}) - 2Cov(G_{1jk}\beta_{dir,k} - G_{2jk}\beta_{dir,k}) + Var(e_{1j} - e_{2j})}$$
$$= \gamma_\Delta^2 \frac{M * \left(\frac{\sigma_d^2}{M} + \frac{\sigma_i^2}{M} + \frac{2\rho_{di}\sigma_d\sigma_i}{M}\right) * \left(2 - 2Cov(G_{1jk}, G_{2jk})\right)}{M * \frac{2\sigma_d^2}{M} * \left(1 - Cov(G_{1jk}, G_{2jk})\right) + (\sigma_e^2 + \sigma_e^2 - 2\rho_e\sigma_e^2)}$$
$$= \gamma_\Delta^2 \frac{2(\sigma_d^2 + \sigma_i^2 + 2\rho_{di}\sigma_d\sigma_i) * \left(1 - Cov(G_{1jk}, G_{2jk})\right)}{2\sigma_d^2\left(1 - Cov(G_{1jk}, G_{2jk})\right) + 2\sigma_e^2(1 - \rho_e)}.$$

When the correlation of genotypes between siblings is 0.5, we can simplify $\gamma_\Delta$ and $R_\Delta^2$ into:

$$\gamma_\Delta = \frac{\sigma_d^2 + \rho_{di}\sigma_d\sigma_i}{\sigma_d^2 + \sigma_i^2 + 2\rho_{di}\sigma_d\sigma_i};$$

$$R_\Delta^2 = \frac{(\sigma_d^2 + \rho_{di}\sigma_d\sigma_i)^2}{(\sigma_d^2 + \sigma_i^2 + 2\rho_{di}\sigma_d\sigma_i) * (\sigma_d^2 + 2\sigma_e^2(1 - \rho_e))}.$$

Further, when the correlation of environmental effect between siblings is 0.5 (i.e., $\rho_e = 0.5$), $R^2_\Delta$ can be simplified as

$$R^2_\Delta = \frac{(\sigma_d^2 + \rho_{di}\sigma_d\sigma_i)^2}{(\sigma_d^2 + \sigma_i^2 + 2\rho_{di}\sigma_d\sigma_i) * (\sigma_d^2 + \sigma_e^2)}.$$

The ratio of $R^2_\Delta$ over the population-level direct effect variance component $h^2{}_{dir,OLS}$ can be derived as

$$\frac{R^2_\Delta}{h^2{}_{dir,OLS}} = \frac{(\sigma_d^2 + \rho_{di}\sigma_d\sigma_i)^2}{(\sigma_d^2 + \sigma_i^2 + 2\rho_{di}\sigma_d\sigma_i) * (\sigma_d^2 + \sigma_e^2)} * \frac{{\sigma_d}^2 + 2{\sigma_i}^2 + 2\rho_{di}\,\sigma_d\sigma_i + {\sigma_e}^2}{{\sigma_d}^2}.$$

When the direct and indirect effects have a correlation of 0 (i.e., $\rho_{di}$), this ratio equals

$$\frac{R^2_\Delta}{h^2{}_{dir,OLS}} = \frac{\sigma_d^2({\sigma_d}^2 + 2{\sigma_i}^2 + {\sigma_e}^2)}{(\sigma_d^2 + \sigma_i^2) * (\sigma_d^2 + \sigma_e^2)};$$

and when the indirect effect is set to zero, this ratio equals

$$\frac{R^2_\Delta}{h^2{}_{dir,OLS}} = \frac{\sigma_d^2({\sigma_d}^2 + {\sigma_e}^2)}{\sigma_d^2(\sigma_d^2 + \sigma_e^2)} = 1.$$

The ratio of $R^2_\Delta$ over the sibling-level direct effect variance component $h^2{}_{dir,\Delta}$ (Appendix 4) can be derived as

$$\frac{R^2_\Delta}{h^2{}_{dir,\Delta}} = \frac{(\sigma_d^2 + \rho_{di}\sigma_d\sigma_i)^2}{(\sigma_d^2 + \sigma_i^2 + 2\rho_{di}\sigma_d\sigma_i) * (\sigma_d^2 + \sigma_e^2)} * \frac{{\sigma_d}^2 + {\sigma_e}^2}{{\sigma_d}^2} = \frac{(\sigma_d^2 + \rho_{di}\sigma_d\sigma_i)^2}{{\sigma_d}^2(\sigma_d^2 + \sigma_i^2 + 2\rho_{di}\sigma_d\sigma_i)}.$$

When the direct and indirect effects are independent (i.e., $\rho_{di} = 0$), this ratio equals

$$\frac{R^2_\Delta}{h^2{}_{dir,\Delta}} = \frac{\sigma_d^2}{\sigma_d^2 + \sigma_i^2}.$$

**4. Within-family regression estimates for direct effect PGS**

Next, we derive the sibling regression results when the direct effect PGS, i.e., $\sum_{k=1}^{M}\big(G_{1jk} - G_{2jk}\big)\beta_{dir,k}$, is known. Regressing sibling differenced phenotype on sibling differenced direct effect PGS, i.e.,

$$Y_{1j} - Y_{2j} = \gamma_{dir,\Delta}\left[\sum_{k=1}^{M}\big(G_{1jk} - G_{2jk}\big)\beta_{dir,k}\right] + e_{dir,\Delta,j}$$

we can get regression coefficient and $R^2$ as follows

$$\begin{aligned}\gamma_{dir,\Delta} &= \frac{Cov\left(PGS_{dir,1j} - PGS_{dir,2j}, Y_{1j} - Y_{2j}\right)}{Var\left(PGS_{dir,1j} - PGS_{dir,2j}\right)} \\ &= \frac{Cov(\sum_{k=1}^{M}\left(G_{1jk} - G_{2jk}\right)\beta_{dir,k}, \sum_{k=1}^{M}\left(G_{1jk} - G_{2jk}\right)\beta_{dir,k} + (e_{1j} - e_{2j}))}{Var\left(\sum_{k=1}^{M}\left(G_{1jk} - G_{2jk}\right)\beta_{dir,k}\right)} \\ &= \frac{Var\left(\sum_{k=1}^{M}\left(G_{1jk} - G_{2jk}\right)\beta_{dir,k}\right)}{Var\left(\sum_{k=1}^{M}\left(G_{1jk} - G_{2jk}\right)\beta_{dir,k}\right)} \\ &= 1;\end{aligned}$$

$$\begin{aligned}R^2_{dir,\Delta} &= \gamma^2_{dir,\Delta} * \frac{Var\left(PGS_{dir,1j} - PGS_{dir,2j}\right)}{Var\left(Y_{1j} - Y_{2j}\right)} \\ &= \gamma^2_{dir,\Delta} * \frac{Var\left(\sum_{k=1}^{M}(G_{1jk} - G_{2jk})\beta_{dir,k}\right)}{Var\left(\sum_{k=1}^{M}(G_{1jk} - G_{2jk})\beta_{dir,k} + \left(e_{1j} - e_{2j}\right)\right)} \\ &= \frac{\sigma_d^2\left(2 - Cov\left(G_{1jk}, G_{2jk}\right)\right)}{\sigma_d^2\left(2 - Cov\left(G_{1jk}, G_{2jk}\right)\right) + 2\sigma_e^2 - 2\rho_e\sigma_e^2}.\end{aligned}$$

When the correlation of sibling genotypes is 0.5, we can simplify this into

$$R^2_{dir,\Delta} = \frac{\sigma_d^2}{\sigma_d^2 + 2\sigma_e^2 - 2\rho_e\sigma_e^2}.$$

When the correlation of environmental effect between siblings is 0.5, this can be further simplified as

$$R^2_{dir,\Delta} = \frac{\sigma_d^2}{\sigma_d^2 + \sigma_e^2}.$$

**5. Rewrite the regression coefficient estimates in Trejo & Domingue (2018)**

Trejo & Domingue constructed their true model for phenotypes and downstream PGS regressions the same way as we did. They focused exclusively on bias in regression coefficient, whereas we focus on $R^2$ and coefficient bias. The setups differ in that Trejo and Dominique "standardize twice"-- they standardized their genotypes and also standardized their downstream PGS once again after all other components in the model were calculated. Therefore, we replicated their calculations except for the "second" standardization in what follows. We use their notation for simplicity.

They denoted their standardized between-family PGS as $\widehat{PGS'}^{D}_{\iota J}$ which equals $\frac{\left(\widehat{PGS}^{D}_{\iota J}-\overline{\widehat{PGS}^{D}_{\iota J}}\right)}{var\left(\widehat{PGS}^{D}_{\iota J}\right)^{\frac{1}{2}}}$, and standardized within-family PGS as $\Delta_0^1\widehat{PGS'}^{D}_{\iota J}$ which equals $\frac{\left(\Delta_0^1\widehat{PGS}^{D}_{\iota J}-\overline{\Delta_0^1\widehat{PGS}^{D}_{\iota J}}\right)}{var\left(\Delta_0^1\widehat{PGS}^{D}_{\iota J}\right)^{\frac{1}{2}}}$. We derived the expression with the non-standardized form, $\widehat{PGS}^{D}_{\iota J}$ and $\Delta_0^1\widehat{PGS}^{D}_{\iota J}$.

Between-family analysis:

$$\hat{\psi}_1 = \frac{cov\left(\widehat{PGS}^{D}_{\iota J}, Y_{ij}\right)}{var\left(\widehat{PGS}^{D}_{\iota J}\right)}$$
$$\hat{\psi}_1 = \frac{cov\left(PGS^{D}_{ij} + PGS^{N}_{ij}, PGS^{D}_{ij} + PGS^{N}_{j}\right)}{var\left(PGS^{D}_{ij} + PGS^{N}_{ij}\right)}$$

Since $PGS^{N}_{ij}$ denotes the transmitted genetic nurture and $PGS^{N}_{j}$ denotes the full genetic nurture from both parents, we can decompose the latter into $PGS^{N}_{ij} + PGS^{NT}_{ij}$, where $PGS^{NT}_{ij}$ denotes the non-transmitted genetic nurture. Since Trejo & Domingue also assumed no assortative mating (thus transmitted and non-transmitted alleles being independent), $cov\left(PGS^{NT}_{ij}, PGS^{D}_{ij}\right) = cov\left(PGS^{NT}_{ij}, PGS^{N}_{ij}\right) = 0$. Therefore, we have

$$\hat{\psi}_1 = \frac{cov\left(PGS^{D}_{ij} + PGS^{N}_{ij}, PGS^{D}_{ij} + PGS^{NT}_{ij} + PGS^{N}_{ij}\right)}{var\left(PGS^{D}_{ij} + PGS^{N}_{ij}\right)}$$
$$\hat{\psi}_1 = \frac{var\left(PGS^{D}_{ij} + PGS^{N}_{ij}\right) + cov\left(PGS^{N}_{ij}, PGS^{NT}_{ij}\right)}{var\left(PGS^{D}_{ij} + PGS^{N}_{ij}\right)}$$
$$\hat{\psi}_1 = \frac{var\left(PGS^{D}_{ij} + PGS^{N}_{ij}\right)}{var\left(PGS^{D}_{ij} + PGS^{N}_{ij}\right)} = 1$$

$$E\left[\hat{\psi}_1\right] = 1$$

Within-family analysis:

$$\hat{\pi}_1 = \frac{cov\left(\Delta_0^1\widehat{PGS}^{D}_{\iota J}, \Delta_0^1 Y_{ij}\right)}{var\left(\Delta_0^1\widehat{PGS}^{D}_{\iota J}\right)}$$
$$\hat{\pi}_1 = \frac{cov\left(\Delta_0^1\widehat{PGS}^{D}_{\iota J}, \Delta_0^1 PGS^{D}_{ij} + \Delta_0^1 PGS^{N}_{j}\right)}{var\left(\Delta_0^1\widehat{PGS}^{D}_{\iota J}\right)}$$
$$\hat{\pi}_1 = \frac{cov\left(\Delta_0^1\widehat{PGS}^{D}_{\iota J}, \Delta_0^1 PGS^{D}_{ij}\right)}{var\left(\Delta_0^1\widehat{PGS}^{D}_{\iota J}\right)}$$

$$\hat{\pi}_1 = \frac{var\left(\Delta_0^1 PGS_{ij}^D\right) + cov\left(\Delta_0^1 PGS_{ij}^N, \Delta_0^1 PGS_{ij}^D\right)}{var\left(\Delta_0^1 PGS_{ij}^D + \Delta_0^1 PGS_{ij}^N\right)}$$

$$E[\hat{\pi}_1] = \frac{1 + \frac{\lambda \rho_g}{2}}{1 + \lambda \rho_g + \frac{\lambda^2}{4}}$$

Thus, Trejo and Domingue's results are consistent with ours with non-standardized estimated PGS for both between-family and within-family analysis.
